# Glycans modulate lipid binding in Lili-Mip lipocalin protein

**DOI:** 10.1101/2023.03.06.531357

**Authors:** Harini SureshKumar, Rajeswari Appadurai, Anand Srivastava

**Affiliations:** Molecular Biophysics Unit, Indian Institute of Science Bangalore, C. V. Raman Road, Bangalore, Karnataka 560012, India

**Author notes:** Electronic mail.

**Keywords:** **Keywords** Lili-Mip, glycans, lipid binding protein, simulations, network analysis, allostery

## Abstract

The unique viviparous Pacific Beetle cockroaches provide nutrition to their embryo by secreting milk proteins Lili-Mip, which is a lipid-binding glycoprotein that crystallizes in vivo. The resolved in vivo crystal structure of variably glycosylated Lili-Mip shows a classical Lipocalin fold with an eight-stranded antiparallel beta-barrel enclosing a fatty acid. The availability of physiologically unaltered glycoprotein structure makes Lili-Mip a very attractive model system to investigate the role of glycans on protein structure, dynamics, and function. Towards that end, we have employed all-atom molecular dynamics simulations on various glycosylated stages of a bound and free Lili-Mip protein and characterized the impact of glycans and the bound lipid on the dynamics of this glycoconjugate. Our work provides important molecular-level mechanistic insights into the role of glycans in the nutrient storage function of the Lili-Mip protein. Our analyses show that the glycans locally stabilize spatially proximal residues and regulate the low amplitude opening motions of the residues at the entrance of the binding pocket. Glycans, which are located at the portal end of the barrel, also restrict the distal barrel depth and allosterically modulate the lipid dynamics in the barrel. A simple but effective distance-based network analysis of the protein also reveals the role of glycans in the subtle rewiring of residues crucial for determining the barrel depth and lipid orientation.

## I. INTRODUCTION

The Pacific beetle cockroach (*Diploptera punctata*) is a unique case of ovoviviparity in insects, where they give birth to live young ones. The gestation period usually lasts for 63 days, wherein the brood sac distends 4-fold and can accommodate up to a dozen embryos. The morphology of the brood sac also further changes to enable sustained secretions of nutrient rich fluid which is ingested by the developing embryos^1^. This feature is a departure from most insect reproductive morphologies, where nutrition is mainly provided by means of yolk deposition in the egg, primarily aided by an exceptionally tuned synthesis of juvenile hormone (JH)^2^. Hence the emergence of this milk protein is a positive selection in response to the evolution of viviparity^3^. As a result of nutritive enrichment, the offsprings are born as advanced instar larvae, which attain reproductive maturity in 43-52 days. The growth is quite fast compared to other ovoviviparous species such as *Rhyparobia maderae* and *Lepidochitona cinerea*, which take more than 90 days^4^. This can be attributed to the substantial deposits of nutrient-rich ‘milk’ fluid in the midgut of embryos and its high calorific value. Biochemical assays show that this milk fluid consists of proteins, glycans and fatty acids^1^. At 27% of gestation period, it replaces the yolk as a nutrient source. To meet the continual consumption of milk by embryos, its synthesis is interestingly high and sustained till parturition. This leads to a 600-fold increase in milk protein. Hence the excess protein is stored via crystallisation, which results in highly concentrated and densely packed reserves of protein and its conjugates^4^. Each crystal of milk protein has 3-4 fold higher calorific value^5^ than its mammalian counterparts and hence has immense potential as an alternate source of nutrition for human consumption.

The milk protein is a lipid binding glycoprotein that has evolved from the Lipocalin family, with a relatively new nutritive role^3^. Interestingly this species reports almost 22 different sequences of milk protein with varied states of glycosylation and lipidation. The lipid binding pocket^5^ is a characteristic *β*-barrel structure which bears similarities to extracellular lipid binding proteins (eLPBs). The fatty acids with carbon chain length between 14 and 18 have the propensity to occupy the barrel, with C18 fatty acid being one among the most preferred^6^ though they have limited effect on the thermal stability of protein^7^. The protein also has four potential glycosylation sites occupied by varied degree and composition of glycans but it is found to be readily crystallizable^6^.

Glycans are generally known to confer diversity to proteins in physical properties and functional properties as well^8^. They are covalently-bound sugar complexes located at a consensus sequence in the protein, assembled via one of the most complex modification processes^9^. Ittypically involves a N- or O-glycosidic linkage between the oligosaccharide units and specific residues of nascent proteins. The former being more common, is generally formed with Asparagine in a Asn-X-Thr recognition sequence whereas the latter is usually formed with Serine or Threonine. Glycosylation is a popular choice of post-transalation modification due to the functional heterogeneity it can provide as a consequence of its structural diversity. Unlike the peptide bonds, the glycosidic bonds are flexible, which allows the protein to sample a wider range of conformations than previously observed^10^. Hence, they alter many aspects of the protein such as its stability, function, interactions, assembly, oligomerization states^11, 12^ with a well-documented example being G-protein coupled receptors^13^. One instance of physical enhancement reported in hCD2ad^14^ increases stability and solubility. It can also regulate functional properties like recognition, site-specific interactions, half-life, functional regulation to name a few^15^. The varying degrees of strong and weak interactions across the glycan chain and protein residues, make interactions highly dynamic and non-specific. Occasionally they can also be more specific, like the glycan-based regulation in endocytic uptake of CD59 by alterations in its surface glycan profile^16^. Glycans can also have case-by-case effects as is seen in stability of proteins in general^17^. Human *α*1-acid glycoprotein is a nice example of glycosylation to escape glomerular filtration in kidneys. Glycosylation is not unusual in lipocalin families or storage proteins like Lili-Mip either. A lot of filarial fatty acid and retinol binding proteins (FAR proteins) are glycosylated but remain to be analysed from the perspective of glycans, though they don’t have structural similarities to Lili-Mip^18^.

Lili-Mip’s implied role as food alternative makes it vital to understand the implications of glycans and lipids on Lili-Mip dynamics. This has been utilised in food industry to mitigate food allergenic response with a case-by-case basis of glycosylation^19^ and deglycoyslation^20, 21^ of proteins. However there are fewer studies on glycans on yolk proteins, stored away in crystalline form, since they usually suffer from micro-heterogeneities like other storage proteins. For example, the folding and assembly of insect storage proteins such as arylphorin^22, 23^ and solubility of vitellogenins^24, 25^, are hypothesised to be affected by glycans. Hence the differential role of glycans have led to an unavailability of a standard prescription to follow when characterising a protein. However in the case of Lili-Mip, the large heterogeneous crystals were resolved with high resolution despite being variably glycosylated with the lipid intact in its barrel. A recent study also suggests that the glycans do not interfere with the protein contact responsible for crystallisation^6^. Though the glycan is taken up as a nutrient alongside Lili-Mip and doesn’t interfere with crystal packing, it is unclear whether glycosylation plays a functional role in addition to being a nutrient. Hence, it is important to explore whether glycans affect the structural stability of the protein. By association, the follow up question would be the effect of glycans on the ligand recruitment behaviour of protein.

*The unaltered form of glycoprotein in the crystalline form implies that a computational study on this structure would reflect on the physiological dynamics of the protein as well.* Hence we have employed all-atom molecular dynamics simulations on various glycosylated stages of a bound and free Lili-Mip protein and characterised the impact of glycans and ligands on the dynamics of this glycoconjugate. We have primarily focused on the changes in physico-chemical aspects and geometry of the protein, that affect its dynamics. Modal analysis and correlation studies have been employed to capture differences in low amplitude functional motions from trajectories. The barrel hole profile is extensively analysed to understand the changes in pocket volume. A simple but effective network model generated from its topology to understand residue-wise contributions to dynamics. Subsequently, the contribution of glycans and ligands to the overall protein dynamics has been elucidated.

## II. MATERIALS AND METHODS

### A. Systems under consideration

To obtain coordinates of the protein structure, we chose PDB accession ID of 4NYQ since it is the natively crystallised protein structure (other structures include PDB ID: 5EPQ, 4NYR). The structure is reported to be glycosylated and ligand-bound, weighing around 20.78 KDa. There are four potential glycosylation sites - Asn44, Asn75, Asn88 and Asn154 observed in the protein. With the exception of site Asn75 (since this monoglycosylated site could only be partially modelled in the crystal structure, it is ignored), the rest are modelled with glycan compositions found in the PDB structure. The list of glycan chains and modifications present in the structure is listed in Table 1.

The protein comprises a signature lipid binding *β*-barrel, which is a characteristic of the lipocalin family^3^ as seen in Fig 1. This barrel accommodates lipophilic ligands like oleic or linoleic acid, in its hydrophobic binding pocket. This barrel is followed by an 15 residue stretch of *α*-helix. The loops between the beta-strands that make up the portal^5^ are named as L1, L3, L5 and L7. Lipid enters and exits through this portal. In this work, we have chosen oleic acid as the model ligand. The glycoprotein is found to be primarily hydrophilic, with an average hydrophobicity score of -0.410.

**FIG. 1:**
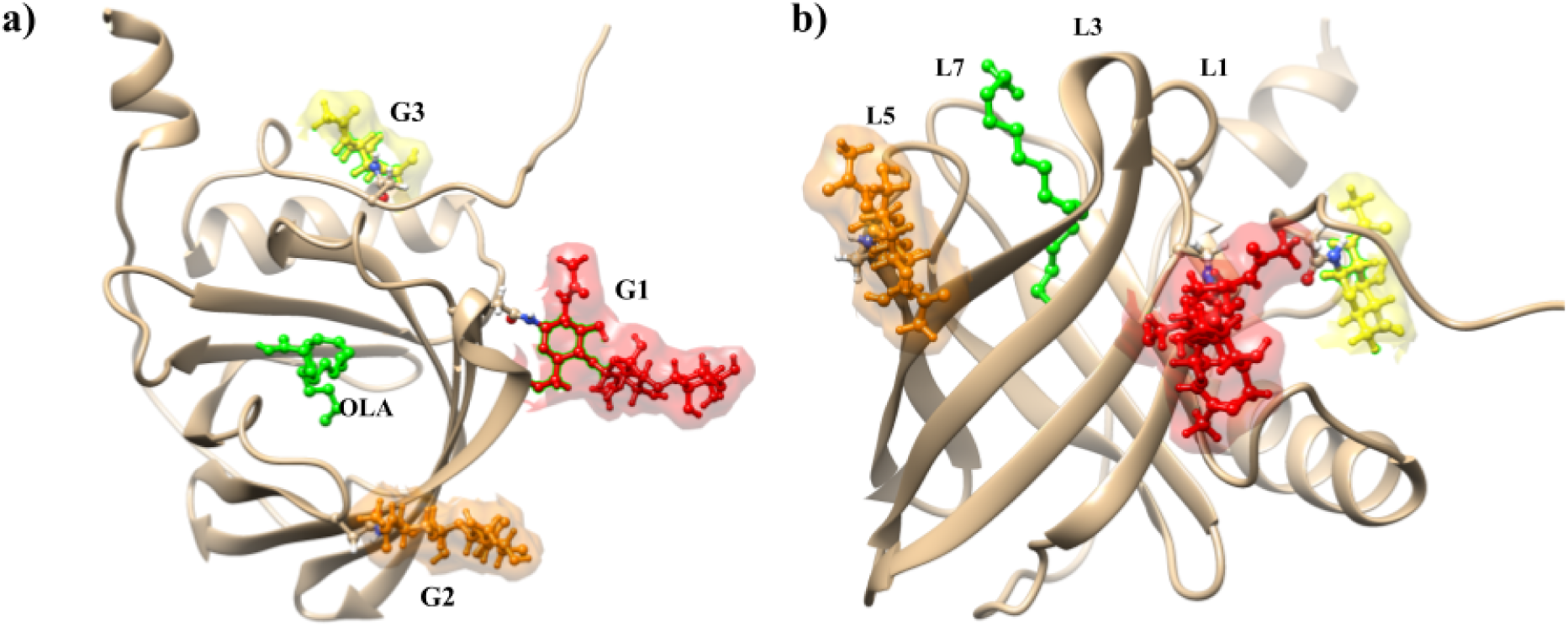
Structure of fully glycosylated and ligand bound Lili-Mip (PDB ID: 4NYQ) a) top view with Oleic acid (green) and glycan chains G1 (red), G2 (orange) and G3 (yellow) b) lateral view highlighting the loops L1, L3, L5 and L7 forming the lipid entry portal

Multiple set-ups were created to study the impact of glycosylation states in the ligand-bound and free protein forms. Firstly to distinguish the role of ligand on protein, we set up lipidated- and delipidated states of a fully glycosylated Lili-Mip (G L and G NL, respectively). To further estimate the individual and co-operative effects of glycans, we constructed mono-glycosylated Lili-Mips and all possible combinations of doubly glycosylated Lili-Mips in its ligand-free state. The set-ups hereafter will be referred to, using the abbreviations listed in Table I.

**TABLE I:** Different set-ups with detailed protein composition for each, is listed here. Each input file consists of the protein in various combinations of glycans and oleic acid

| Set-up | Detailed Composition | Ligand | Glycosylation |
| --- | --- | --- | --- |
| G_L | Fully glycosylated,Ligand bound | yes | full |
| NG_L | Unglycosylated,Ligand bound | yes | no |
| G_NL | Fully glycosylated, Ligand-free | no | full |
| NG_NL | Nascent protein | no | no |
| G1_NL | Glycosylated at 1st site, Ligand-free | no | Only in 1st site |
| G2_NL | Glycosylated at 2nd site, Ligand-free | no | Only in 2nd site |
| G3_NL | Glycosylated at 3rd site, Ligand-free | no | Only in 3rd site |
| G12_NL | Glycosylated at sites 1 and 2, Ligand free | no | At site 1,2 |
| G13_NL | Glycosylated at sites 1 and 3, Ligand free | no | At site 1,3 |
| G23_NL | Glycosylated at sites 2 and 3, Ligand free | no | At site 2,3 |

### B. Molecular Dynamics Simulations

All input and paramete files are prepared using CHARMM-GUI^26–29^. The glycans, lipids, 8 N-terminal residues and 2 C-Terminal residues are modelled using the ‘Glycan reader and modeller’ package of CHARMM-GUI. Each setup is then solvated with TIP3P water model in a cubic box of volume 405 nm^3^ and neutralised with potassium and chloride ions. We then performed Molecular Dynamics (MD) simulations using GROMACS 5.1.4 wherein the interactions were described by CHARMM-36 force field^30, 31^. The set-ups are minimised using steepest descent algorithm in the Verlet cut-off scheme to treat bad contacts. LINCS algorithm restrains the bonds involving hydrogen. PME was utilised for the calculation of long-range electrostatic forces with a cut-off of 12 *Å* for short-ranged interactions. The setups are equilibrated for 125 ps with a time step of 1 fs to achieve convergence in energy and density. Nose-Hoover thermostat and Parinello-Rahman barostat maintains the set-ups at 300 K and 1bar. Non-bonded interactions were limited to an inter-atomic distance of 10 *Å* by the method of force switch. The production run was for 800 ns with a time step of integration set to 2fs.

Input files needed to initiate molecular simulations and full trajectory data of all simulations for all systems considered in this work are available on our server for download. The server data can be accessed via our laboratory GitHub link: codesrivastavalab/Lipocalinglycan-glycan. The full trajectory files for all the systems under consideration can also be accessed directly from our SharePoint location, which is provided in the github link.

### C. Trajectory analysis

#### 1. Analyses of protein dynamics

Trajectory analysis was performed using the analysis modules of GROMACS 5.1.4^32, 33^ and R-Bio3d package^34^. For preliminary analysis, the global conformational changes are investigated using root mean square deviation (RMSD), which is the mean square displacement of protein conformation evolving with time. In addition to RMSD, the detailed residue-level fluctuations are also investigated using root mean square fluctuations (RMSF), which is averaged across time for each residue. An R-based package, Bio3D, was utilised for further structural analysis. Further, principal component analysis (PCA) was employed to capture the slow modes that resolve the essential cooperative motions in the protein. Dynamic cross correlation (DCCM) was assessed for each residue pair (j, k) to understand residue-level correlations.

#### 2. Barrel pocket volume analysis

The changes in barrel radii are calculated using HOLE program suite of MDAnalysis 3.2.^35, 36^. The profiles are generated every 5 ns and the barrel radii in each profile are calculated along the barrel axis. The radii slices start from 50 *Å* and end at 30.2 *Å* of the barrel height along its axis. The slices are then assembled to reconstruct the barrel profile. Since the exact opening of the barrel is uncertain due to fluctuations, the calculations exclude the topmost regions of the portal which are highly dynamic and statistically poor to infer from.

#### 3. Network analyses

A distance-based protein structure network was generated and analysed using NetworkX suite of Python^37^. To achieve a dynamic network system, we have constructed network every 100 ps for the last 700 ns, that tracks conformational changes in the protein. Each residue is reduced to a node and are considered to be linked if any of the inter-residue atomic distances fall within 7 *Å*. Such a liberal cut-off ensures that the more dynamic counterparts of the protein might be partially linked. The distances are now supplied as edge weights to each edge. They are immediately reweighed to discard redundant edges in the network and amplify the influences of nodes in the network. The reweighing is done using edge betweenness centrality, where the edges are weighed based on its occurrence in all shortest paths in the network. This ensures that the edges central to communication flow are scored higher in the network. In addition, the redundant paths arising due to larger distance cut-off are also taken care of.

We then utilise centrality measures to capture residue-level changes in the network. Centrality captures the hotspots or hubs that are central to structural preservation and/or information flow within the network. We utilised node based metrics that are sensitive to conformational changes in the protein. Eigenvector centrality (EVC) scores nodes based on its contact strength and the influence of its neighbouring residues. Hence, if a node is located in a densely connected area, its centrality will be high. This is complemented by the characteristic path length (ΔCPL) analysis, which is a path-dependent metric and is hence sensitive to conformational changes in the protein. It measures the change in average path length of the network after the removal of node and its associated edges. Higher ΔCPL scores indicate the importance of a particular amino acid for information flow in the network. We further calculated ΔΔCPL calculation between glycosylated and deglycosylated systems to understand the effect of glycan on the ΔCPL score of amino acids.

## III. RESULT AND DISCUSSION

We have discussed the results in four sections. Firstly, we investigate the impact of glycans on protein structure and its dynamics. Secondly, we explore the role of glycans on barrel architecture and lipid dynamics. Lastly we deduce how the post-translational modification by the addition of glycans that occurs several Angstrom away from the lipid binding site, communicate/couple to the structurally and functionally critical barrel residues and to the lipid substrate using protein structural network analyses.

### A. Global structure of Lili-Mip is preserved but glycans induce functionally relevant fluctuations locally

We simulate the native Lili-mip protein monomer consisting of covalently attached glycans and non-covalently bound lipid substrate (GL system) in solution. The initial structure obtained from the PDB equilibrated using molecular simulations and exhibited expected fluctuations. The initial structural relaxation data is presented in Fig. S1. After this initial structural relaxation, further change in structure occur more gradually indicating equilibrium sampling in all the four systems including the G NL and NG NL where we removed the lipid substrate, indicating that the structural integrity is maintained even in the absence of ligand that conforms to previous findings^7^. Comparing all the 4 systems we observed that the glycosylated forms (G L and G NL) have relatively larger values of RMSD than the non-glycosylated forms. In order to deconvolute the effect of individual glycans we then simulated Lili-Mip after systematically removing the glycans one after the other. We observed that the largest structural deviation in protein is essentially caused by the presence of glycan-2 when it is co-occurring with either of the other glycans. Individually the glycans 1, 2 and 3 have lesser and almost similar RMSD vales (Fig S1.b).

Next, we measured the RMSF to see the residue wise fluctuations and presented the data for the major 4 systems in Fig. 2.a and the other systems in Fig S1.d. As the figures indicate, the changes inherent to the protein mostly arise from residues forming the terminus and lipid portal comprised of loops L1, L3, L5 and L7 with the largest fluctuation occurring around loop 3. The proximity of glycans 1 and 2 further increases the flexibility of L3 in G L and G NL systems. In conjunction with this, the exclusion of considering the flexible loops for the RMSD measurements reduces the value to the sub-Angstrom level in the non glycosylated systems indicating that the barrel architecture is quite rigid without glycans. However, in presence of glycans the RMSD did not quench only by the exclusion of loops but also required to eliminate certain deeper beta barrel residues (Fig. 2.b, Fig S1.c) flanking the portal loops (residues 63, 64, 69, 70, 71, 72, 73, 86, 87, 95 and 96). These analyses suggest that the glycans impart greater flexibility particularly at residues near the lipid binding portal loops. But it is to be noted that none of these effects lead to collapse or structural destabilization of protein since radius of gyration values (Fig S2) suggest good compaction in all cases^38^ indicating global integrity.

**FIG. 2:**
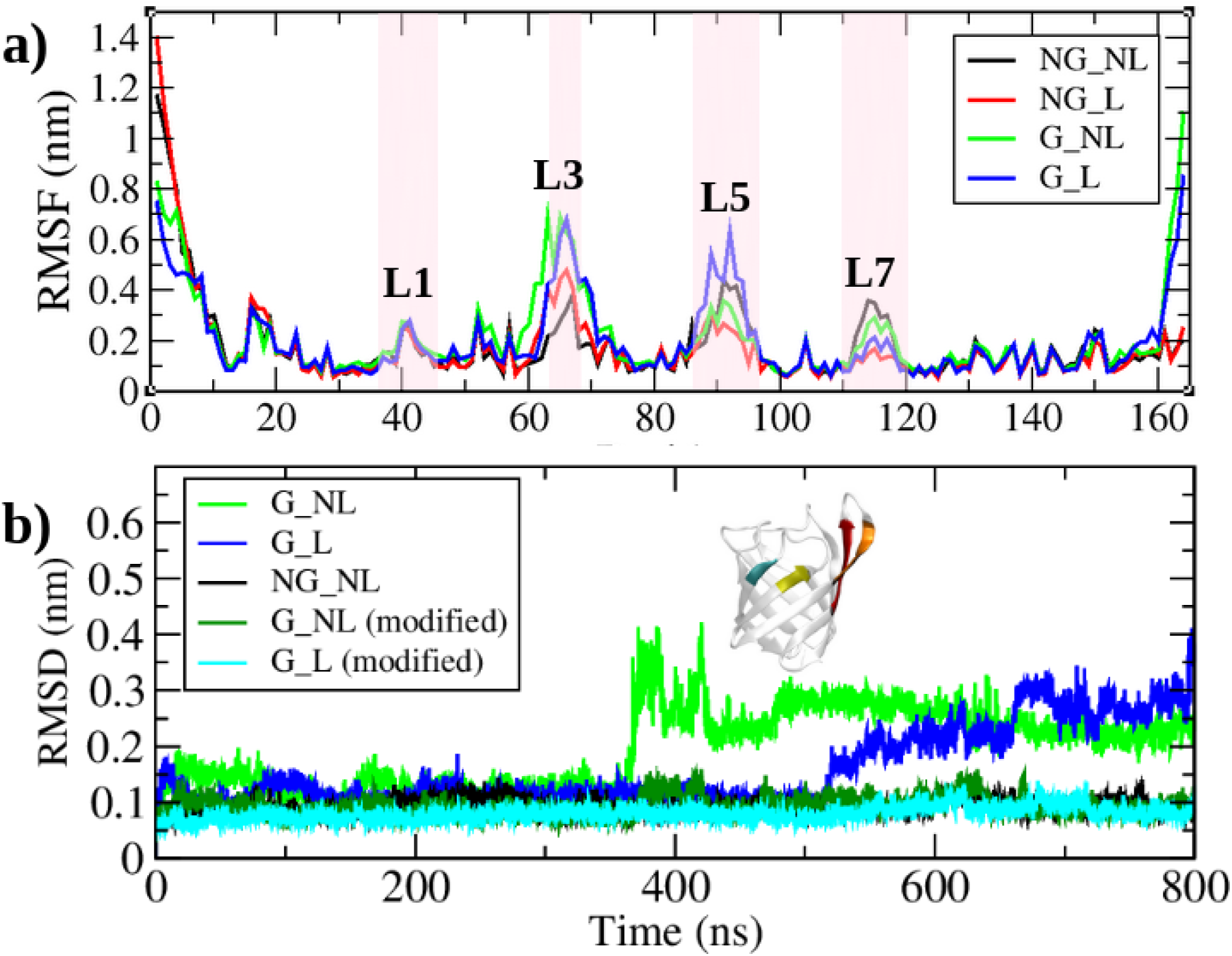
(a) RMSF of all four lipocalin set-ups (b) RMSD of C-*α* carbons of secondary structures in glycosylated systems G NL and G L. RMSD is recalculated (modified) by removing residues 59-64 (red), 69-73 (orange), 86-87 (yellow), 95-96 (teal) from G NL (dark green) and G L (cyan) setups

The dynamics of the loops however, are markedly different in glycosylated and deglycosylated Lili-Mip in delipidated form. The collective motions of the loops are identified by time-averaged Pearson’s correlations (Fig 3). The cross-correlation map thus constructed, is first validated by the secondary structure signatures occurring along the off-diagonals (Fig S3). From the outset, there are large stretches of protein that undergo collective motions (positively correlated) in the native state. Upon further scrutiny, we see that the correlations between loops shift when incorporating glycans into the system. Glycans cause strong negative correlations amongst loops, regardless of the presence of ligand. The primary motion in glycosylated Lili-Mip is captured by the slowest mode of principal component analysis.

**FIG. 3:**
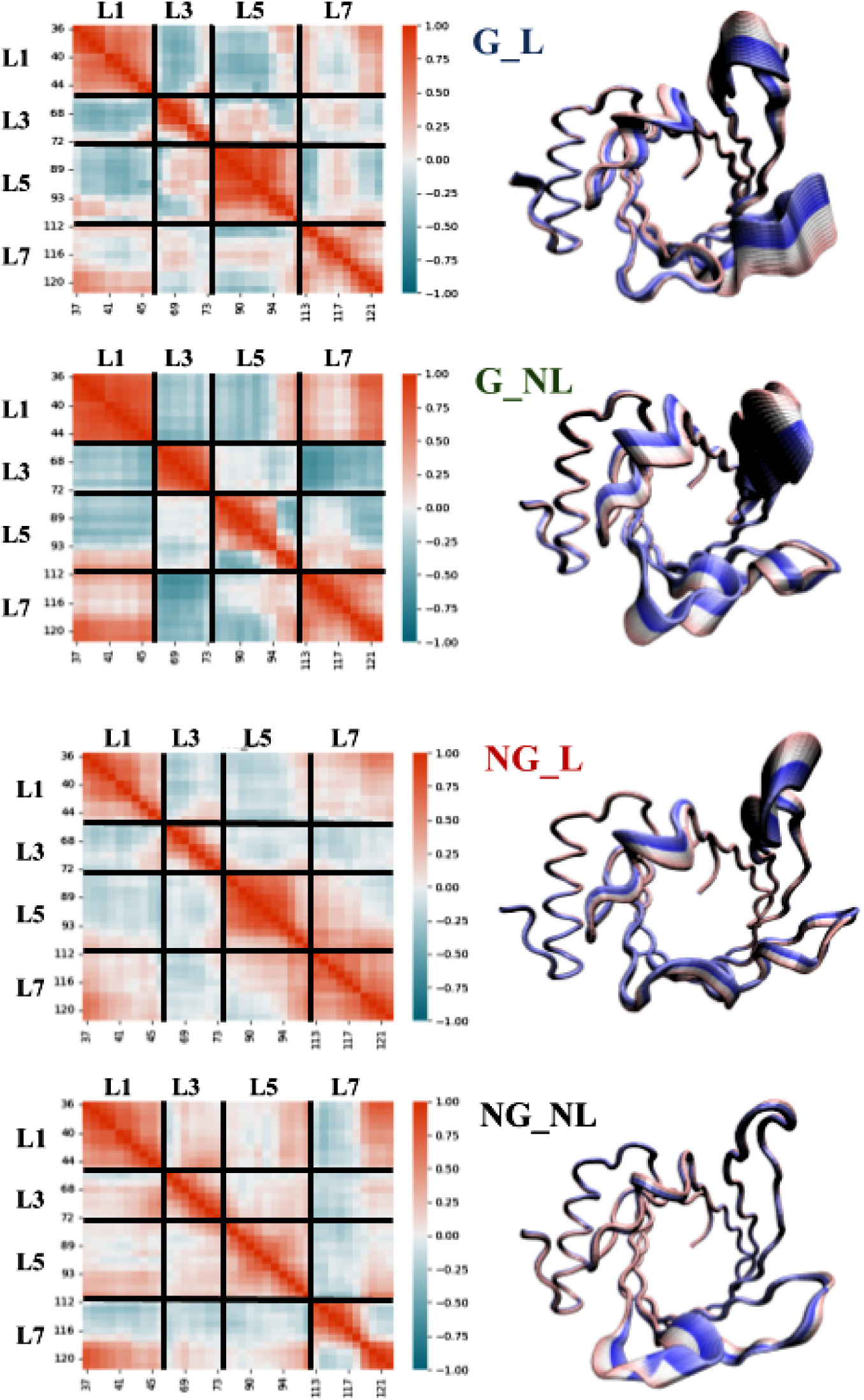
Dynamic cross correlation at the loops of portal in Lili-Mip in the primary four systems. A positive correlation is indicated by red and negative correlation is indicated by Blue. The slowest mode of the protein (from principal component analysis (PCA) is represented for all four major systems beside their respective correlation maps. The evolving positions of the loops with respect to time, are color coded from pink to dark blue. We show the mode analyses based dynamical motions for all four systems in movie file movieS1.mp4 in the SI

The loops function as a single entity which result in an opening-closing motion at the portal, which might be involved in the lipid recruitment process. The pairs L3-L7 and L1-L5 are clearly anti-correlated which allows breathing motion to take place. On the other hand, the deglycosylated form, though exhibiting fluctuations, is weakly correlated at the portal. Only loop L7 is seen to be partially anti-correlated with other loops. The deglycosylated Lili-Mip is rather rigid. Together, we structural and dynamical analyses show that glycans introduce functionally relevant and fine-tuned localised motions in Lili-Mip. The absence of glycan subdues the portal movements, and the portal size of delipidated Lili-Mip is subsequently stabilised.

### B. A complete ligand re-orientation takes place in Lili-Mip due to the glycans

The correlations between lipid and residues however are completely different (Fig 4.a). The ligand, oleic acid, has distinct correlation patterns with certain residues (loops L1, L2, beta-strands 1,2 and 3). This set is weakly anti-correlated with OLA patch1 (carbons C1 to C8 in oleic acid) in the glycosylated system. However, in deglycosylated Lili-Mip, the same set of residues are now anti-correlated with OLA patch2 (carbons C9 till C18 in oleic acid). Moreover the overall correlations are much weaker in the deglycosylated system, which suggests either decorrelated motion or stabilisation of oleic acid fluctuation profile. Hence we examined the dynamics of oleic acid in the lipidated system G L and NG L.

**FIG. 4:**
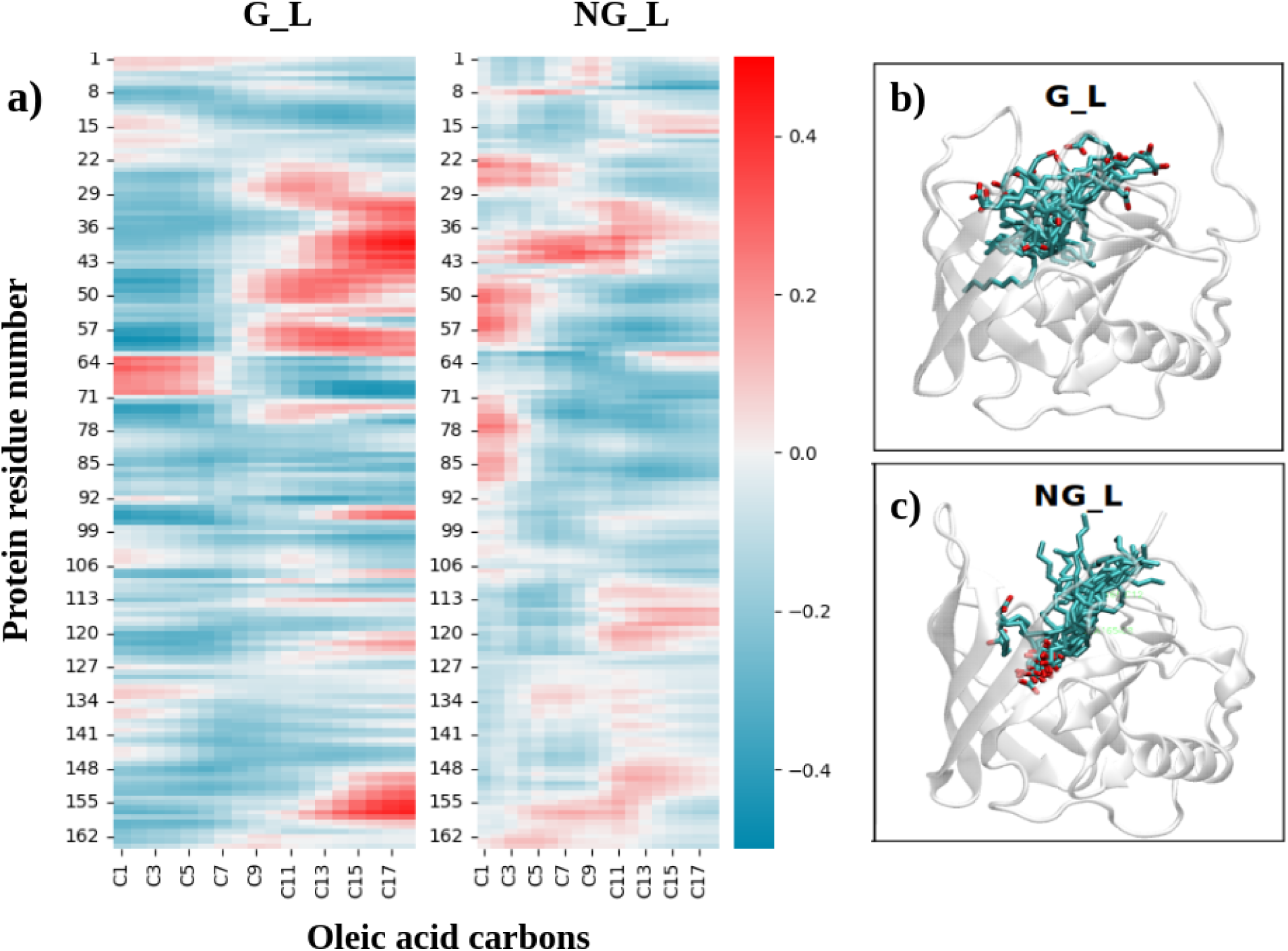
(a) Cross correlation map between C1-C20 of Oleic acid (OLA) and amino acids inLili-Mip. Oleic acid re-orientation profile superposed every 50 ns for (b) G L set-up and (c) NG L setup. Headgroup is distinguished by the oxygen of carboxylate group colored in red

It was surprising to note that these two systems have completely different orientations. In glycosylated Lili-Mip (Fig 4.b), the lipid exhibits conformational flexibility but is mostly confined to the portal surface. The carboxylate group is solvent-exposed which is also orientation reported during structural characterization of this protein^5, 6^. However in deglycosylated Lili-Mip (Fig 4.c), the lipid is tethered headfirst in the barrel, parallel to its axis. The surface accessibility of oleic acid in deglycosylated Lili-Mip also drops down by 110 *Å*^2^ approximately (Fig 5.a). The barrel architecture must undergo considerable re-arrangements to accommodate these changes, apart from the flexibility imparted by glycans. This is discussed in detailed in upcoming section.

**FIG. 5:**
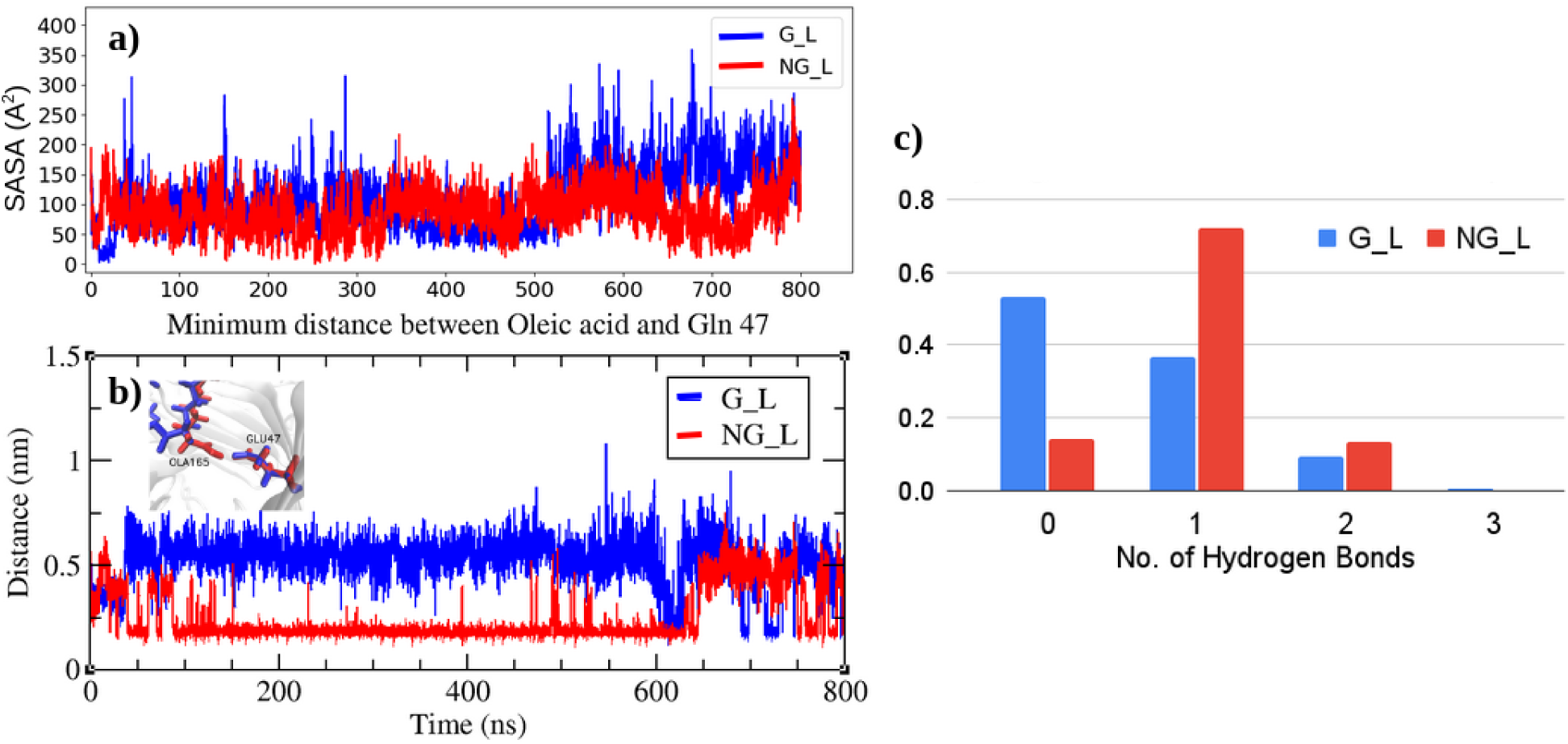
(a) Solvent accessibility surface area (SASA) reported for oleic acid in glycosylated (G L) and deglycosylated (NG L) set-ups (b) Depth accessibility of oleic acid by measuring the minimal distance between E47 and oleic acid. Inset shows the distance between E47 and OLA in G NL (red) and G L (blue) set-ups (c) Normalised histogram of all hydrogen bonds formed between Oleic acid and residues for G L and NG L set-ups

Lipid orientations aren’t uncommon and have been observed in most intracellular lipid binding proteins (iLBPs)^39^. The proteins have an inward oriented lipid which form tight hydrogen bonds with conserved residues, and sometimes also through ordered water molecules. For example, Liver fatty acid binding protein (LFABP) is found to accommodate two lipids in different orientations whereas intracellular lipid binding proteins like retinol binding protein, accommodate lipid in inward orientation. Proteins like MFB1 whose residue composition and orientation aid the barrel shape rearrangement, can accommodate lipids in a slightly different orientation, where it is almost perpendicular to the barrel axis^39^. The binding affinities are enthalpic in nature and mostly in the micro-molar range. However this holds true at physiological temperatures with relative binding affinities for different fatty acids^40^. Especially in the case of L-FABP, the differences in binding affinities of oleate at both the sites vary by almost 20 fold^41, 42^. There are even reports of lipocalins (olfactory binding proteins) whose glycosylation states dictate the ligand specificity^43^. This leads to low-affinity and high-affinity binding sites, wherein the former is located closer to the portal, with a conformationally flexible bound-lipid. Hence the barrel shape and the fine-tuning of residue orientations might play an important role in lipid binding, orientation and lipid type specificity^44^. Consequently, we proceeded to analyse differences in contact between barrel residues and oleic acid.

We posit that there must be strong interactions that secure the lipid in inward orientation in deglycosylated Lili-Mip. We observed that one particular residue Glu47, which is buried deep in the pocket, was in close contact, around 1.9 *Å* (Fig 5.b) with the ligand. But they are nearly 6 A apart in the native structure. There is also a sustained and specific hydrogen bonding formed between certain residues (Glu47, Asp62, Thr90, Asp89) and oleic acid that anchor the lipid in this form (Fig 5.b inset). However the distribution of hydrogen bonds are spread out (Fig 5.c)and promiscuous indicating non-specific hydrogen bonds formed at the surface, causing a weak retention of lipid at the surface. Since none of these residues are spatially proximal to glycans, there might be an allosteric regulation taking place.

At this point we must be cognizant of the fact that such transitions in lipid orientations can take place only if the barrel structure is sufficiently flexible. However former studies have shown that the glycans do not affect the overall pocket volume of Lili-Mip^7^. Hence a subtle re-arrangement of interior residue orientations might facilitate the lipid’s access to the buried residues. We hence profiled the barrel along its axis to understand the architectural changes due to the incorporation of glycans and ligand, which we will discussing in the next section.

### C. Glycans allosterically modulate the barrel profile

To study the barrel profile in detail, we calculated the barrel radii at different depths along the barrel axis at every timestep. The barrel shape is re-constructed in Fig 6 with the slices of mean barrel radii stacked upon each other. The dashed lines represent maximal standard deviation from the mean radius. The major part of the barrel of lipidated Lili-Mip set-ups are gourd shaped whereas the delipidated-Lili-Mip profiles (inset figures) resemble aconical funnel.

**FIG. 6:**
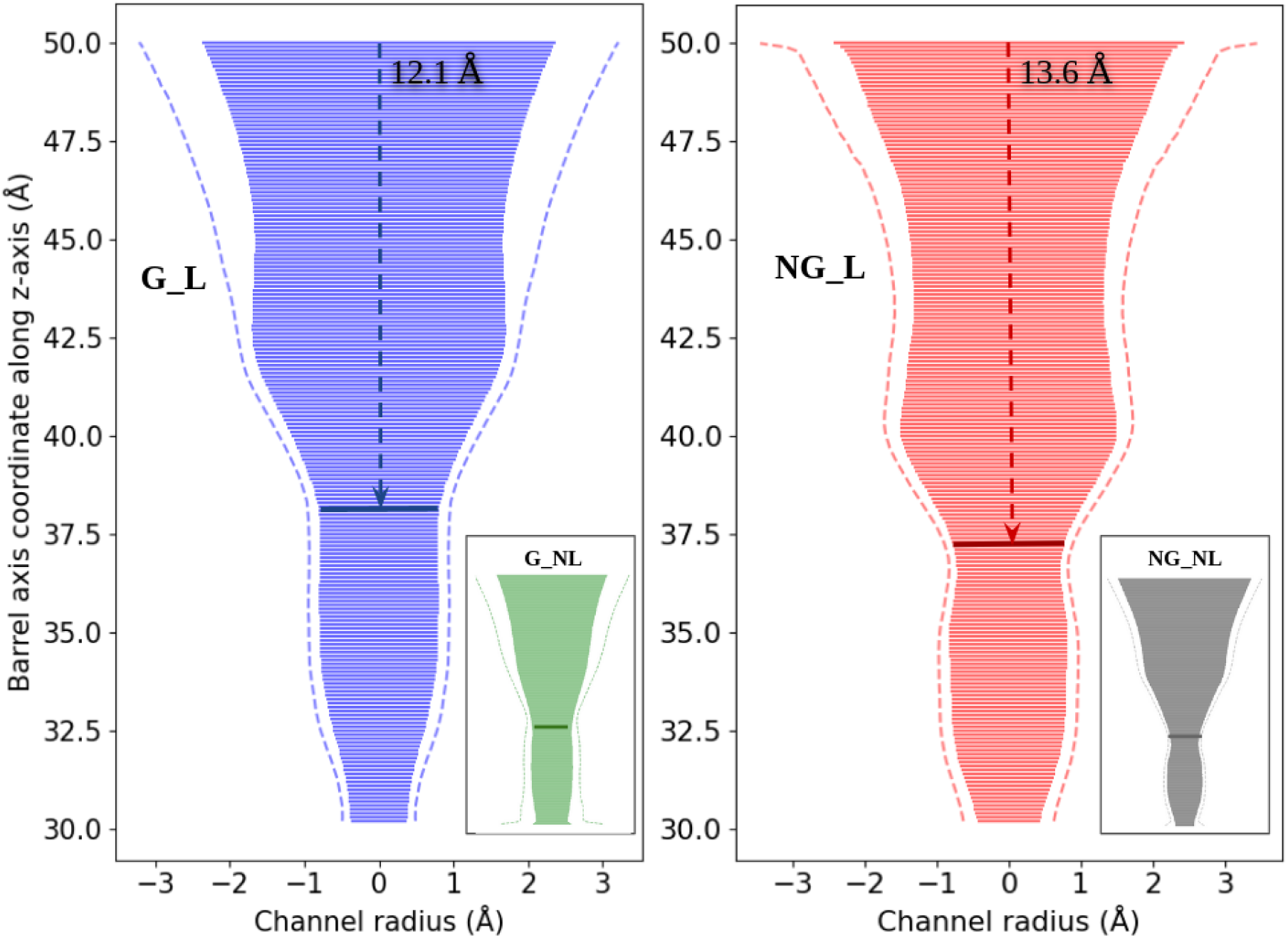
Barrel profile in solid color, with its maximum possible standard deviation (in dotted lines) reported for lipidated set-ups (G L in blue and NG L in red). The delipidated counterparts (G NL in green and NG NL in grey)are in the insets. Barrel depth is denoted in Angstroms. The depth limit is marked by a solid line around 37.5 *Å* for all set-ups

We first studied the barrel opening radii that serves as lipid entry portal. The high standard deviation in the glycosylated system profile alludes to the increased flexibility of the backbone in barrel residues. When plotted in a time-dependent manner, the portal radii fluctuates sinusoidally. It must be noted that despite the induced flexibility by glycans, the set-up with largest opening is deglycosylated delipidated-state (NG NL), which explains the motions from principal component analyses. Though the portal of glycosylated Lili-Mip is smaller than NG NL, it is flexible enough to attain sizes comparable to NG NL’s portal.. Hence glycans actually reduce the barrel opening significantly, but the localised fluctuations still allows large portal openings. This is also observed in the PCA images where the overall barrel opening size is reduced in glycosylated set-ups (Fig 3). Hence NG NL might be the most preferred state for lipid recruitment, due to its portal radius.

Next we analysed the profiles of delipidated systems, followed by lipidated systems. Glycosylated Lili-Mip profile in delipidated state (inset-green) resembles an hour-glass. The constriction in the barrel reduces the depth of accessibility ligand but is sufficient to let awater molecule pass through. Upon closer inspection, we observe the twist in *β*-strands, broaden the funnel radii at the top. But the interactions of external residues such as 61 and 69, are stabilised quickly. It also explains the change in RMSD of well-ordered regions of protein without affecting the barrel compaction. This behaviour is absent in deglycosylated Lili-Mip, cementing the notion that glycans cause the *β*-strand twist. Hence the deviations in profile is also less. Absence of glycans to induce flexibility near portal end of barrel, allows a slightly deeper access to the pocket (around 0.6 Å), which is otherwise unavailable.

The lipidated system profiles are markedly different from that of delipidated set-ups. Firstly the presence of lipid stabilises the *β*-strands that were quite flexible in delipidated-Lili-Mip. Hence the profile deviations are much lesser in both glycosylation states. Occupancy of lipid, causes distension mid-barrel, giving rise to a peculiar gourd shape. The first inflection point of the gourd-like profile might be the determining factor that allows lipid orientation, which we will shortly explain in detail. The second inflection point value corresponds to the depth delimiting segment, which is slightly above the radius of water probe (1.6 Å). When measured till this point, we observe that the depth is restricted in case of glycosylated Lili-Mip by almost 1.5 Å. The portal radii in NG L (4.7 Å) and G L (4.9 Å) are similar and hence do not affect lipid recruitment or release. However the distended region in G L is closer to the surface by almost 1.5 *Å* in which the lipid is found to explore multiple conformations. This is also due to the transient hydrogen bonds it forms (see Fig 5) with multiple residues from N terminus (residues 1 till 5) and *β*-strands (Ile38, Lys40, Tyr93, Pro114, Ser118). Hence a barrel depth difference of 1.5 *Å* in addition to the elevation of the binding region in G L, can dictate the lipid type, since the protein is designed to chaperone a variety of lipids to the embryo mid-gut. Ramaswamy’s group established that the residue Glu47, along with Phe85, Phe109, Tyr93, Tyr97 and Leu122 form the depth delimiters of the barrel that subsequently determine the lipid size^5^. From the barrel profiles, we deduce that the determining factor might be the barrel free volume made available by the glycans. As a consequence, certain internal residues (Glu47 (buried), Asp62 and Tyr93) are available to form sustained hydrogen bonds with oleic acid which determines its binding mode. Hence many residues act in concert to change the barrel profile, which in-turn dictates the lipid binding modes.

### D. Network analyses capture the subtle rewiring of residues crucial for determining the barrel depth and lipid orientation mediated by Glycan

Changes in the barrel profile must be facilitated by re-arrangement of contacts in the system, to permit flexibility in lipid orientation. Rather than investigating the modification from elaborate time-dependent distance profiles, we opt for large scale network analysis, to compare the residue level features between the proteins in presence and absence of glycans, in both lipidated and delipidated conditions. The protein topology is first reduced to nodes and edges, every 100 ps. We treat amino acids as nodes and link them with edges, if any of their heavy atoms are within a distance of 7 *Å*. The edges are weighted based on the distances such that the spatially closer residues are strongly-connected (or more weighted) than the other counterparts. Further, to emphasize the cost of information flow, which occurs generally through the shortest path, we re-weighted the edges by the edge-betweenness centrality. This measure scores edges based on its occurrence in all shortest paths in the network and thus highlights the optimal paths while discarding the multiple redundant ones in the network.

We then assess the level of influence of each residue (node) for the network communications using two independent node centrality scores. *Higher scores generally belong to conserved residues that are central to information flow during conformational changes.* The first measure, eigenvector centrality (EVC), is based on degree (number of connections) of a node and its neighbors. It is a nuanced metric that distinguishes equally well-connected residues and hence is sensitive to geometrical rearrangements in the network. We normalised the scores for every network and reported the mean normalised scores for each residue. We provide the above data as a CSV file (network-analysis.csv) in the SI). We assigned importance only to residues whose scores lay in the 4th quartile of their score distribution. Such well-connected residues are observed to be localised at the mid-axis of the barrel, that connects the helix, in all set-ups of the protein (Fig 7.a). A few residues lose influence upon lipidation as expected. However in glycosylated set-up, the locality is narrowed down to an arc along the mid-axis comprising specific residues - Thr34, Leu83, Ile99, Val102, Phe109, Leu120, His124. However, the formation of denser connections at localized regions of proteins *may or may not correspond directly to the information transfer*.

**FIG. 7:**
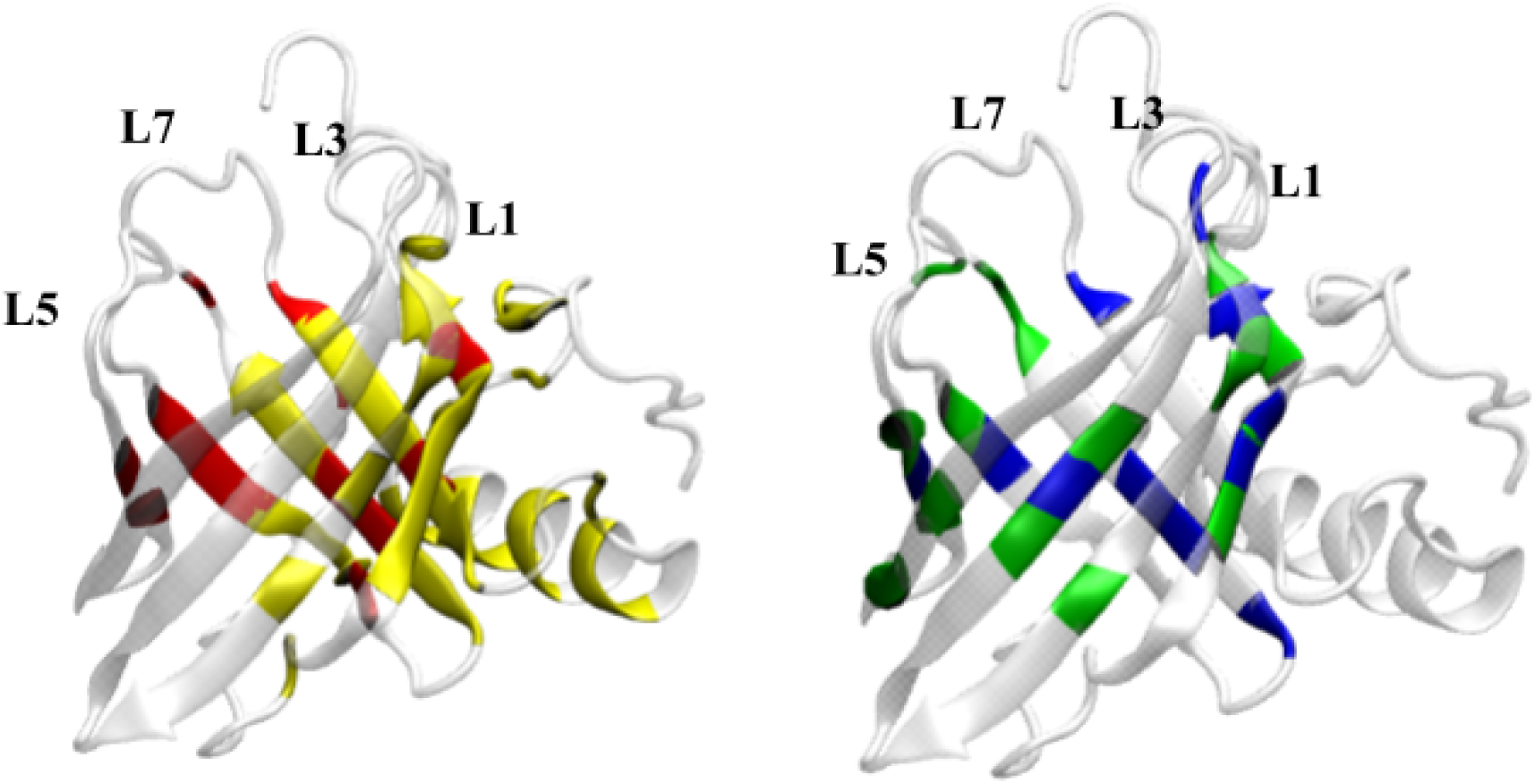
a) Residues with consistently high z scores in Eigenvector centrality in all 4 systems are in yellow. The residues with highest score in ΔΔCPL due to glycans are in red. An arc is formed by red coded residues along the barrel mid-axis b) Residues with highest changes in path length (ΔΔ CPL) when incorporating glycans (G L - NG L) in lipidated-Lili-Mip (Blue) and incorporating glycans (G NL - NG NL) in delipidated-Lili-Mip (Green)

Hence to check that if the residues act as switch for the essential communications inthe network, we then measured variation in characteristic path length (ΔCPL), where we find changes in network’s average path length upon removal of a target residue and its corresponding edges. A residue is deemed important for information flow, if its absence increases the CPL score. Hence in our case a residue with positive ΔCPL (CPL(i) - CPL *>* 0) is influential. We normalised the scores for every frame and considered only positive values lying in the 4th quartile of the score distribution, for our study. Immediately, we see that the set of influential residues or ’hotspots’ are preserved across all protein set-ups. However, their extent of influence differs at different systems. For instance, the presence of glycan changes the network in both the lipidated and delipidated proteins. We bring out that by measuring the ΔΔCPL in both lipidated (ΔΔCPL (L)= G L - NG L) and delipidated systems (ΔΔCPL (NL)= G NL - NG NL). We again consider only the scores from the 4th quartile of the score distribution for our analysis.

Higher ΔΔCPL scores will indicate that the these residues now serve as channels of transmission during a conformational change, in presence of glycans. Such residues are highlighted appropriately in Fig. 7 for lipidated- (blue) and delipidated (green) set-Lili-Mip set-ups. An interesting pattern in glycosylated set-up of delipidated form is the formation of two modular rings - one located above the mid-axis of the barrel, closer to the portal and another towards the distal end of the barrel. This is similar to the residues in arc from EVC measure, albeit a complete structure. The residues making up this arc, from the internal region of the barrel are Leu31, Thr34, Val42, Ile45, Tyr49, Val60, Val74, Val78, Tyr97, Phe109, Pro113, Leu120. Since the protein is more dynamic in presence of glycan, more number of residues are required to sustain the architecture of the barrel. Most changes are through side chain rotations (refer Fig S5), whereas some are influenced by backbone dynamics of the residues. For example, phenylalanine residues 97 and 109 protrude into the barrel space of NG NL set-up. However, their rings are more parallel to the barrel axis, upon glycosylation. Valine (Val60) scores high only in G NL system and contributes to reduction in barrel radius. This is purely influenced by backbone dynamics, in absence of ligand, which allows it to come in contact with Phe109. Upon lipidation, the b-turn is disallowed, which also renders this amino acid redundant.

In a ligand binding protein such as Lili-Mip, the ligand usually functions as external structural stabilizing agent that re-centralise the communication pathways around them. We observed this from the ΔCPL score of ligand and the ΔCPL score distribution of residues. Also, the distribution is less spread in lipidated systems, with multiple residues losing its influence on pathways formerly important in delipidated Lili-Mip. The set of residues now central to the communication pathways in the set-up are now exactly at the barrel mid-axis and only form one ring comprising internal residues Leu31, Thr34, Val42, Ile45, Val60, Phe85, Phe109 and Leu120. This section also corresponds to the depth delimiter region and the score captures 3 of the residues (Phe85, Phe109 and Leu120) responsible for the restriction. Further visual observations show that the glycosylated set-up has small co-operative motions in barrel internal residues, that result in protrusion of side chains into the cavity (Fig S4). Similar to the conditions of aromatic rings observed in delipidated system, the rings are seen to be significantly protruberant in glycosylated form. Residues Phe85 and Phe109 are also closer in contact by 1 *Å* as a consequence, which leads to the depth restriction. Hence most barrel profile re-arrangements are due to cooperative side chain re-orientations influenced by backbone dynamics, which inturn accommodates lipid flexibility in glycosylated systems.

Lastly, we compared the influence of lipids and glycans on each other. The EVC indicates that the glycans significantly lower the influence of lipid (ΔΔEVC = -0.4) whereas it doesn’t affect the communication pathways set by the lipid (ΔΔCPL = -0.04). However presence of lipid consistently elevates the importance of Glycan G3 as a contact and a hub. This is possibly due to its location between the helix-barrel interface. When the barrel is further opened upon lipidation, G3 being the sole external additive, might anchor the interactions at the interface. This observation is further supported by the general increase in CPL and EVC score of barrel residues near glycan G3. However G2 and G3 are affected conversely. It is to be noted that unlike G3, the general ΔCPL scores of glycans G2 and G3 are well-below the mean, in addition to them being in the periphery of the network. Since the applied metrics are sensitive to the geometry of the network, the effect of peripheral nodes might not reliably estimated. However the significant differences in scores allows us to speculate a possible hierarchy of lipid or glycan occupation in the protein. The lipidation has minimal effect on glycan G1’s neighbourhood influence in the network. However G2 has a negative effect on the protein communication pathways and contacts, since it destabilises b-strands 4 and 5. The effect on G2 is vastly improved only upon lipidation by anchoring this section. G1 does not display drastic changes in the metric scores, but the trends are similar upon lipidation. Though the contacts are more or less unperturbed, lipidating a glycosylated protein leads to multiple re-arrangement of pathways. This indicates that the glycosylated protein must undergo major conformational changes when accommodating the lipid. However glycosylating the lipidated form of protein requires far less re-arrangement of pathways in the network. Hence it might be favorable for the protein to be lipidated prior to glycosylation. This is further strengthened by the fact that the barrel radii is the largest in the nascent protein state (NG NL). Hence glycan is not essential for recruitment of lipid. However it induces co-operative opening and closing motions at the portal and restricts the barrel depth, which in turn can determine the specificity of the lipid. Glycosylated protein also retains the lipid in a solvent-exposed orientation which may lead to better dissociation. Hence glycans may actually be involved in the expulsion of the lipid, thus making it easily available for nutrient consumption.

## IV. CONCLUSION

Lili-Mip has been shown to provide a rich nutrient supply to developing embryos of Pacific cockroaches, consisting of proteins, sugar and lipids. It has been dubbed as a super-food with a potential for human consumption. The nutritive milk Lili-Mip is a lipid-binding glycoprotein that can accommodate different lipids in a non-specific manner.The glycosylation is meant to be an addition that serves nutritive purposes primarily. However a molecular dynamics study performed on Lili-Mip protein of the lipocalin-binding family has shed light on the effect of glycosylation in protein dynamics, and lipid recruitment as a consequence. This protein is seen to be conformationally stable in all conditions in the in-cellulo conditions. However the presence of glycans serve as more than just additives. The glycans in no linear fashion, cause conformational changes in the protein. They create highly tuned local but functionally relevant fluctuations in the protein. Glycan 1 and 2 are primary targets hence due modification can result in a structurally stabler form of protein. The resulting residue-level fluctuations are collective in nature and lead to opening-closing motion of the portal. Interestingly we find that the portal opening is highest in its deglycosylated form which might be more favorable for lipid entry, albeit showcasing rigidity in its motions. However oleic acid binds in two different modes, depending on the glycosylation state of the protein. This difference arises due to differences in barrel geometries which determine the binding partners for the fatty acid. The intended effect is the restriction of barrel depth, upon lipidation. There is an allosteric effect of glycans at play, that leads to rearrangement of barrel interior architecture. We lastly employed network analyses to pinpoint the barrel residues that are affected by glycans in lipidated and delipidated conditions. Hence mutating these residues might lead to conformational and functional instability. On the corollary, the unimportant residues can also be mutated, for nutrition enhancement. Though glycans do not alter the contacts, they have a non-linear effect on Lili-Mip dynamics. hence their composition and location becomes important. A secondary inference was the order of assembly of this milk protein. Correlation studies and PCA have suggested that the correlation map and motion of a lipidated system is closer to that of a native structure. The portal opening of deglycosylated state is the largest as well. Hence the deglycosylated state might actually be favorable for lipid recruitment. However we deduce that a more transient lipid binding is preferred for the purposes of quick entry and release during consumption. The glycosylated form retains this ligand binding mode. Network analysis also suggests that the addition of glycans after lipidation, do not induce major conformational switches in the protein. Hence we speculate that the protein might recruitment lipid first, to stabilise its barrel, followed by glycosylation, which now favors the release of lipid from its binding pocket. Hence glycans have shown to have an extended function determining beyond its nutritive role in milk protein.

## Supporting information

network analysis file

Mode Analyses dynamical motions

## V. AUTHOR CONTRIBUTIONS

AS conceived the idea and formulated the project with help of HS. HS constructed and set up models, carried out simulations and generated all the data, figures and plots for the manuscripts. HS, RA and AS analyzed the data and wrote the paper together.

## ACKNOWLEDGMENTS

We would like to thank Ramray Bhat, Purusharth Rajyaguru, Ramaswamy Subramanian and Partha Radhakrishnan Santhakumari for their valuable inputs. HS would like to acknowledge financial support from Ministry of Education, Government of India. R.A. would like to thank the early career fellowship from the DBT-Wellcome Trust Indian Alliance. A.S. thanks the early career grant from the Department of Science and Technology (DST) of India. A.S. also thanks the DST for the National Supercomputing Mission grant. Financial support from the Indian Institute of Science-Bangalore and the high-performance computing facility ”Beagle” setup from grants by a partnership between the Department of Biotechnology of India and the Indian Institute of Science (IISc-DBT partnership programme) are greatly acknowledged. FIST program sponsored by the DST and and UGC, Centre for Advanced Studies and Ministry of Human Resource Development, India is gratefully acknowledged by the authors.

## I. TABULATED DATA

### A. Heteroatom composition

The composition of heteroatoms namely glycan chains and ligand are briefed in the Table S1 listed below. Special bonds such as disulfide bonds are also mentioned but are not investigated in this study. However its role has been linked to regulating the pH-dependent solubilisation of crystals**^?^** .

**TABLE I:** List of non-protein molecules in the protein Lili-Mip

|  |  |
| --- | --- |
| Glycans | β-D' GlcNAc (1 → 4) β-D' GlcNAc (1 →)-44<br>β-D' GlcNAc (1 → 4) β-D' GlcNAc (1 →)-88<br>β-D' GlcNAc (1 →)-154 |
| Ligand | Oleic acid |
| Di-sulphide bridges | 13 ( → ) 146<br>129 ( → ) 160 |

### B. Hydrogen bonds formed between residues and lipid

The non-bonded interactions established between glycans and ligand, oleic acid with the residues of the protein are deduced using VMD Hydrogen bond analysis module. The list of residues maintaining hydrogen bond with ligand and glycans vary significantly over the trajectory. The details are listed below in these tabulations .

**TABLE II:** List of residues forming hydrogen bonds with ligand

| Set-up | Residues forming hydrogen bonds with ligand |
| --- | --- |
| G_L | Ala2, Ala3, Leu5, Asp37, Glu47, Asp62, Tyr93, Pro114, Ser118, |
| NG_L | Glu47, Asp62, Thr90, Asp89, Tyr49 |

### C. Barrel depth analysis

The inflections in the barrel profile occur at different points in each system. For standardization we have measured the inflection at barrel width constrictions 3.5 A and 1.5 A. The depths measured at these sites are listed below

**TABLE III:** Barrel at different inflection points

| System | First inflection | Second inflection |
| --- | --- | --- |
| G_L | 3.5 Å at 42.5 Å | 1.5 Å at 37.9 Å |
| NG_L | 3 Å at 40.03 Å | 1.5 Å at 36.4 Å |
| G_NL | - | 1.5 Å at 37.5 Å |
| NG_NL | - | 1.5 Å at 36.8 Å |

The hydrogen bonds between residues and glycans are briefed in different set-ups, which emphasizes the impact of different glycan environments.

**TABLE IV:** List of residues forming hydrogen bonds with glycans

| Glycans | Set-up | Residues forming Hydrogen bonds with glycans |
| --- | --- | --- |
| G1 | G_L | 43, 64, 157, 32, 66, 156, 162, 128, 46 |
|  | G_NL | 43, 157, 66, 70, 69, 46, 164, 67, 63 |
|  | G1_NL | 164, 28, 32, 161, 163, 157, 162, 46, 159, 30, 61, 63 |
|  | G12_NL | 43, 157, 46, 61 |
|  | G13_NL | 157, 43, 46, 32, 28, 164, 30 |
| G2 | G_L | 89, 52, 75, 70, 73, 86, 57, 71, 87, 86 |
|  | G_NL | 75, 89, 57, 73, 86, 88 |
|  | G2_NL | 89, 73, 75, 57, 55, 86, 72, 52 |
|  | G12_NL | 70, 75, 73, 89, 71, 72, 90, 52, 57, 59, 64 |
|  | G23_NL | 89, 73, 75, 55, 57, 62, 72, 54, 86 |
| G3 | G_L | 152, 150, 149, 162, 163, 164, 125 |
|  | G_NL | 152, 150, 164, 149, 150, 152, 162 |
|  | G3_NL | 152, 150, 163, 149, 164, 125, 162, 161 |
|  | G13_NL | 152, 164, 150, 162, 163, 149, 161 |
|  | G23_NL | 152, 150, 164, 163, 149, 162, 161, 125 |

### D. Network analysis

The residues scoring Z-scores greater than 0.5 in Eigenvector centrality (Table S6) and in Characteristic path length (Δ CPL) centrality (Table S7), of the protein are listed here. The residues gaining importance after the glycans are incorporated (ΔΔCPL) are picked out by difference (Table S8) in G L and NG L system (blue) incase of lipidated systems and in similar fashion for delipidated systems (green). The ubiquitously important residues are highlighted in darkgreen.

**TABLE V:** Residues with high z-scores in Eigenvector centrality

| Z score | Colour | Residues |
| --- | --- | --- |
| 1-1.5 | yellow | 29, 32, 34, 39, 101, 106, 120, 130, 149 |
| 1.5-2.0 | orange | 31, 33, 35, 100, 110, 121, 138, 142, 147 |
| Above 2 | red | 107, 108, 109, 122, 123, 124, 125, 152 |

**TABLE VI:** Residues with high z-scores in ΔCPL

| Z score | Colour | Residues |
| --- | --- | --- |
| 1-1.5 | yellow | 1, 32, 45, 76, 100, 108, 120, 123 |
| 1.5-2.0 | orange | 25, 31, 66, 67, 83, 110, 125, 126 |
| Above 2 | red | 29, 47, 85, 97, 107, 109, 122, 124 |

**TABLE VII:** Residues recording high changes in CPL after glycan is added to lipidated and delipidated Lili-Mip (ΔΔCPL)

| Z score | Colour | Residues |
| --- | --- | --- |
| Lipidated | blue | 32, 34, 35, 47, 85, 98, 106, 108, 120, 123 |
| Delipidated | green | 45, 61, 65, 74, 78, 82, 86, 93, 97, 108, 113, 120, 125 |
| Common residues | dark green | 17, 30, 31, 42, 60, 109 |

## II. FIGURES

**FIG. 1:**
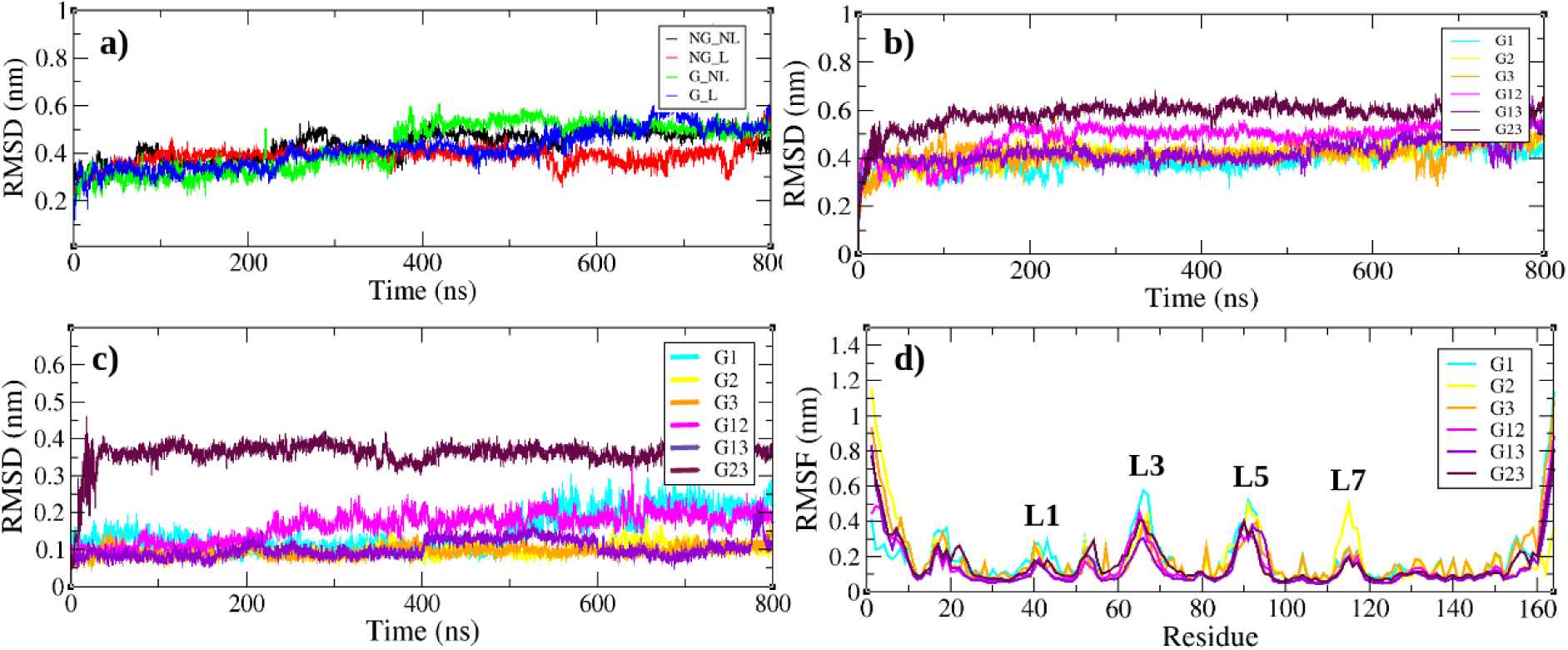
(a) RMSD of C-*α* carbons of whole protein Lili-Mip (b) RMSD of C-*α* carbons of secondary structure residues of glycosylated states of apo-Lili-Mip (c) RMSF of C-*α* carbons of all residues at different glycosylation levels of apo-Lili-Mip

**FIG. II:**
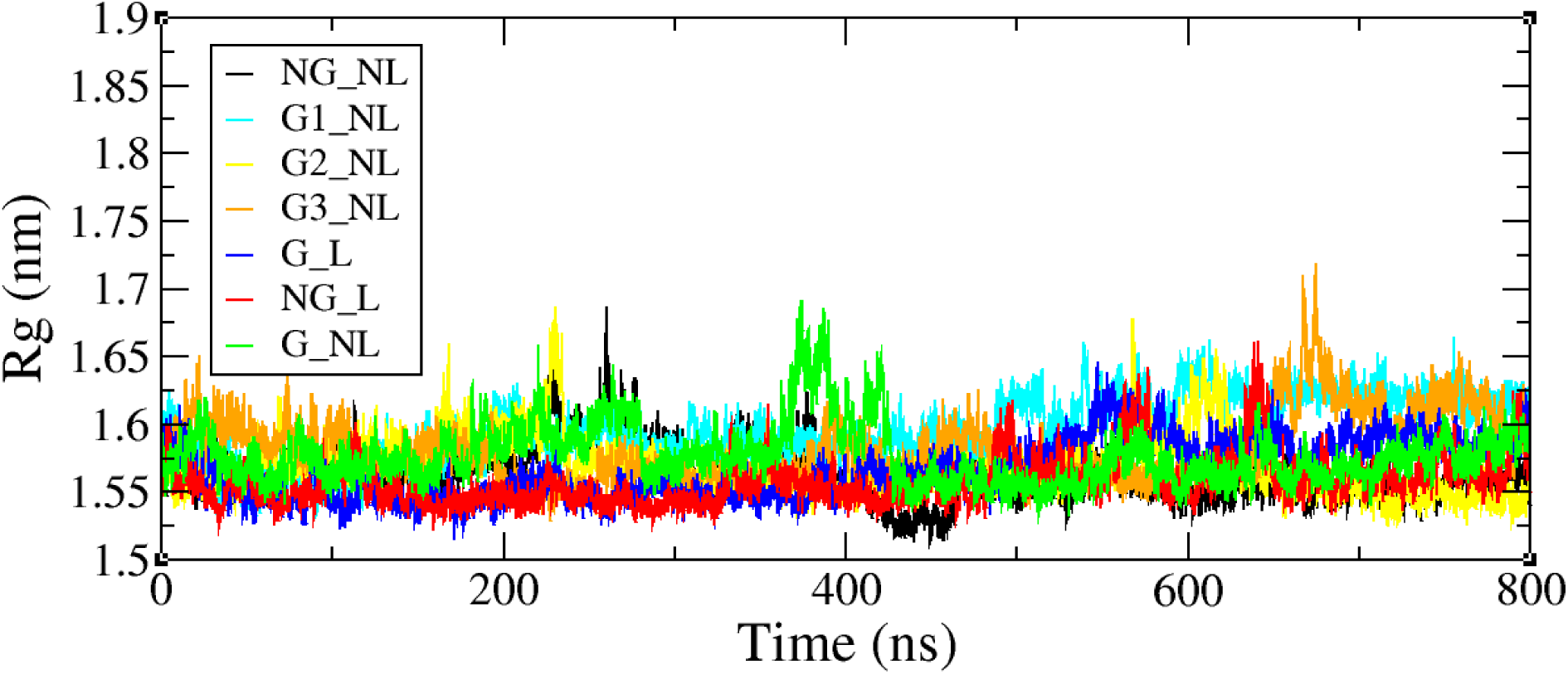
Radius of Gyration of Lili-Mip in different set-ups. The values are similar in most cases except glycoyslated systems G3 NL and G1 NL

**FIG. III:**
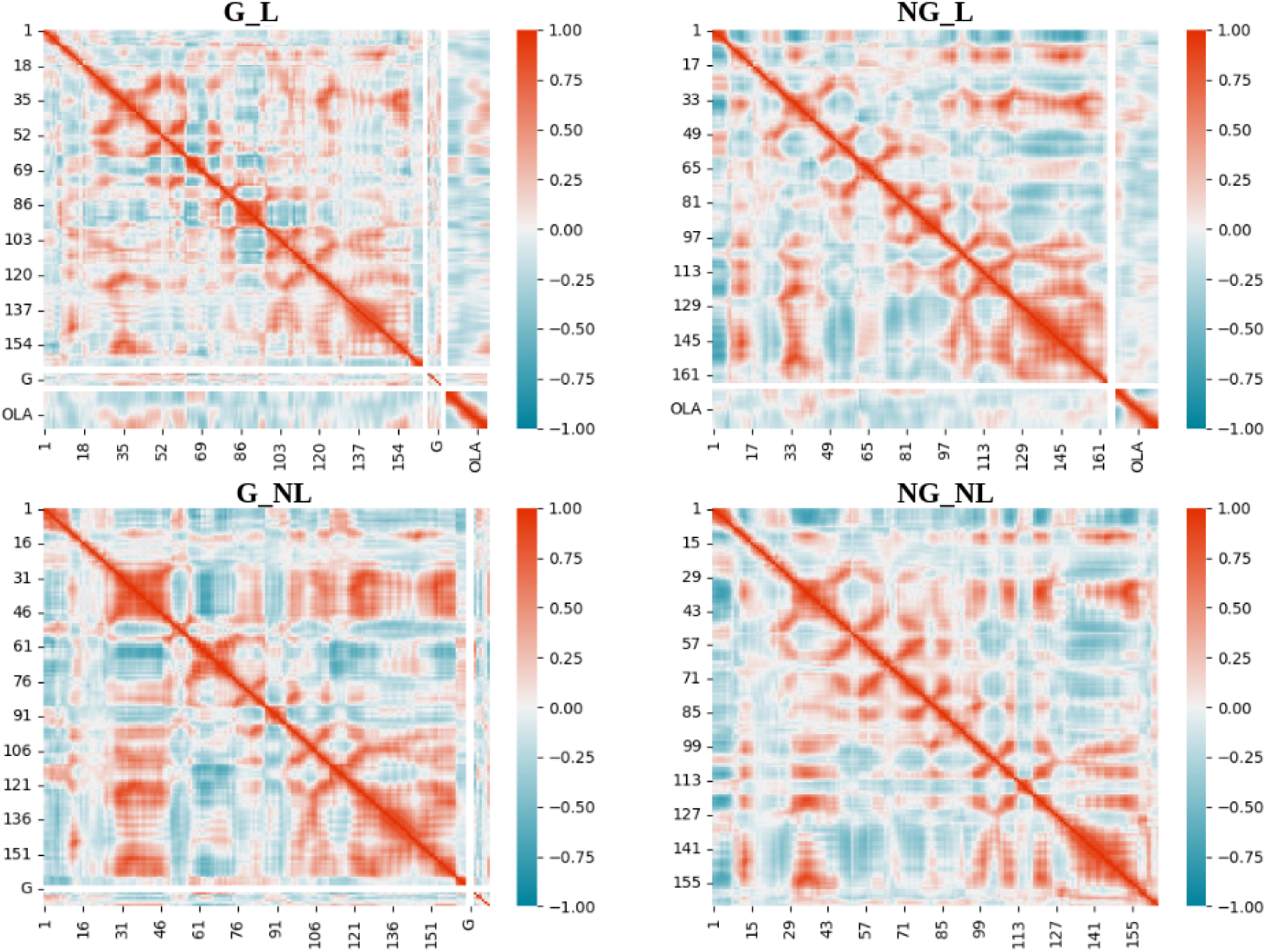
Dynamic cross correlation of residue level fluctuations in the first four primary set-ups. Positive correlations (red) and negative correlations (blue) range from 1 to -1 respectively

**FIG. IV:**
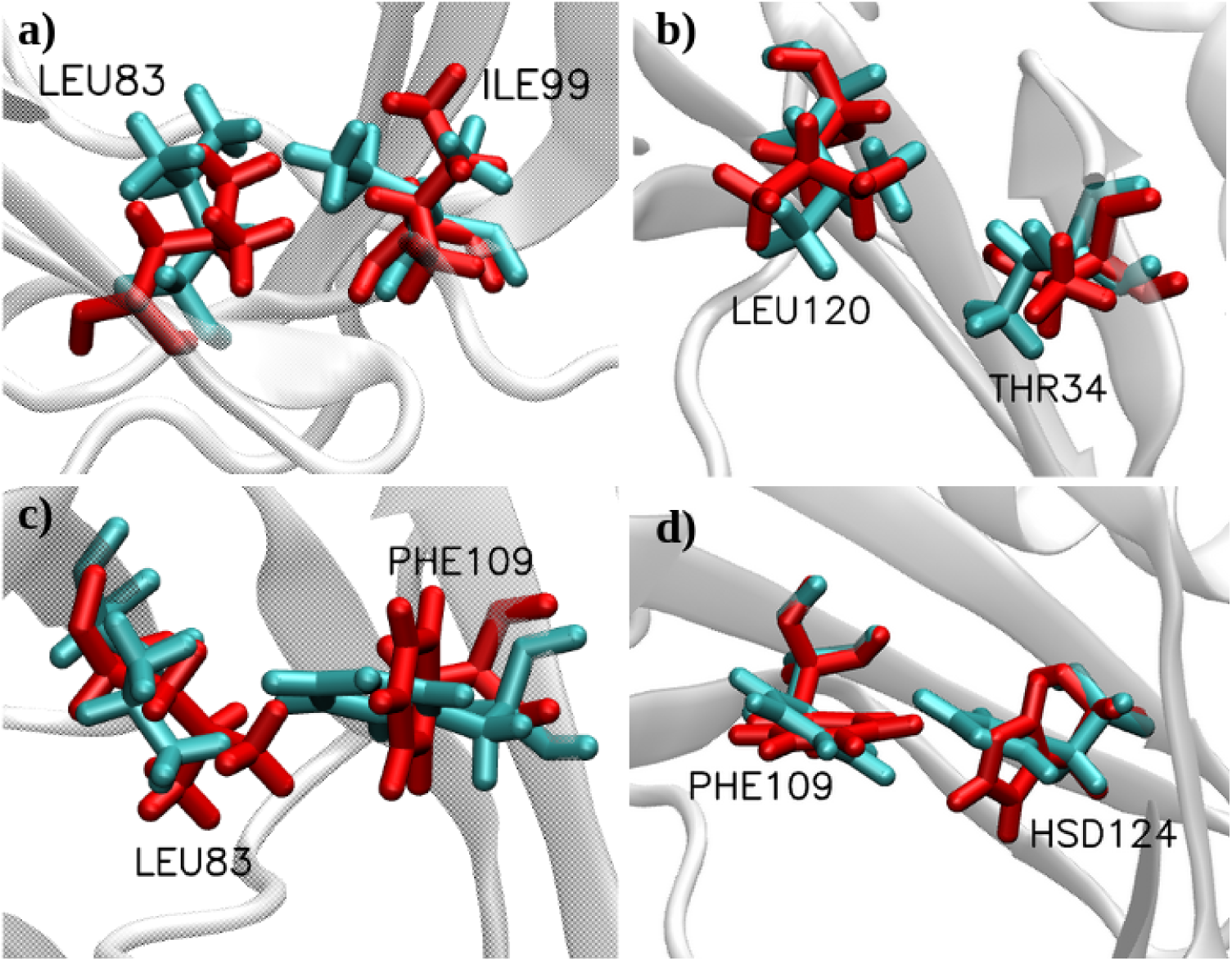
Individual snapshots of side chain orientations of aminoacids are displayed for glycoyslated (blue) and deglycosylated (red) apo-Lili-Mip. In all cases the side chains are closely associated for glycosylated systems.

**FIG. V:**
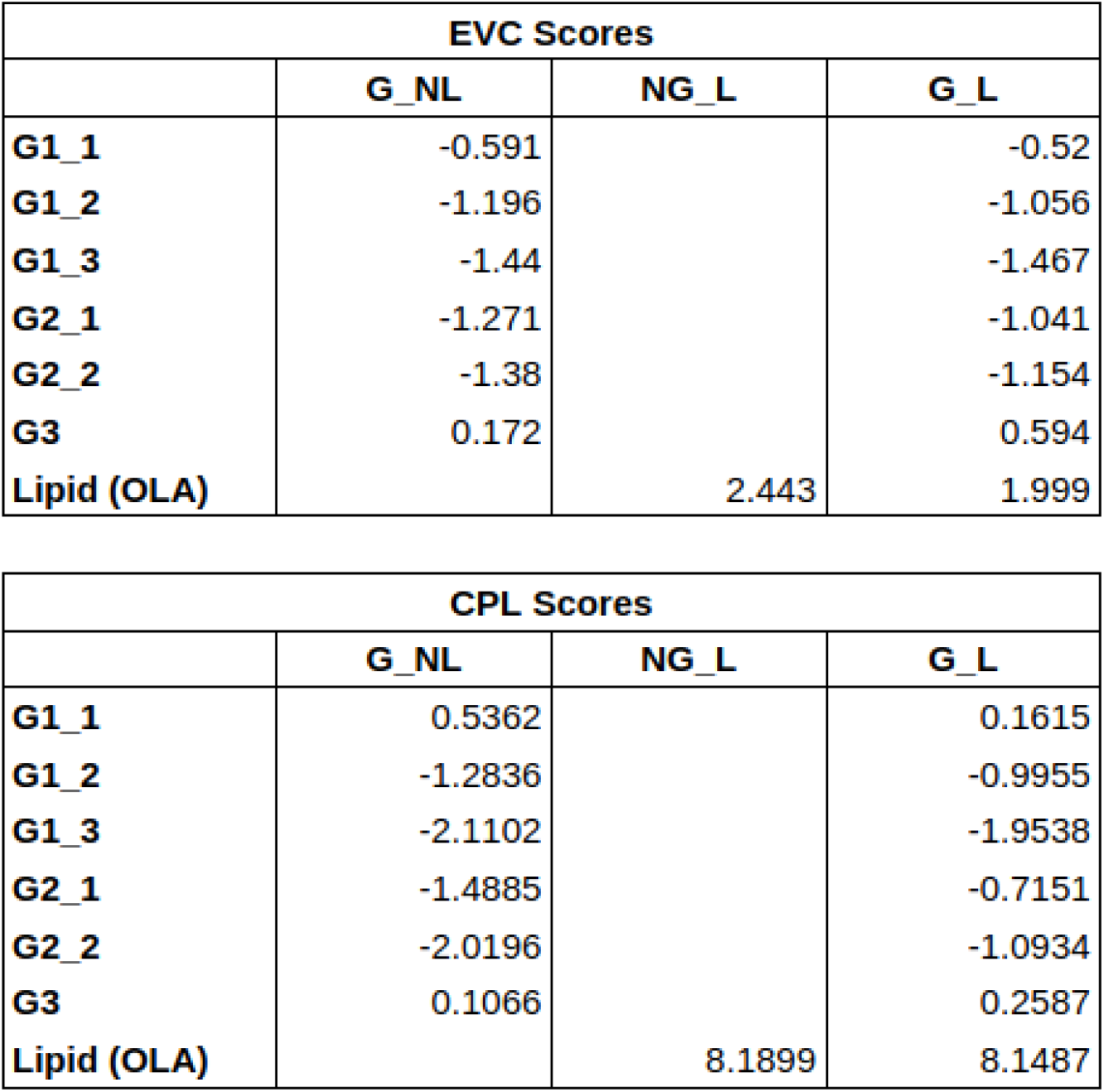
EVC and Δ CPL scores for glycans and lipid in the four primary setups. The ΔCPL scores greater than 0.5 are color coded in shades of green in the set-ups. The changes in ΔCPL of glycans when introducing lipids (G L - G NL) and changes in ΔCPL of lipids when introducing glycans (G L - NG L) are highlighted in color gradations according to their differences

